# Subject-specific whole-brain parcellations of nodes and boundaries are modulated differently under 10Hz rTMS

**DOI:** 10.1101/2021.03.09.434571

**Authors:** Vladimir Belov, Vladislav Kozyrev, Aditya Singh, Matthew D. Sacchet, Roberto Goya-Maldonado

## Abstract

Repetitive transcranial magnetic stimulation (rTMS) has gained considerable importance in the treatment of neuropsychiatric disorders, including major depression. However, it is not yet understood how rTMS alters brain’s functional connectivity. Here we report changes in functional connectivity captured by resting state functional magnetic resonance imaging (rsfMRI) within the first hour after 10Hz rTMS. We apply subject-specific parcellation schemes to detect changes (1) in network nodes, where the strongest functional connectivity of regions is observed, and (2) in network boundaries, where functional transitions between regions occur. We use support vector machines (SVM), a widely used machine learning algorithm that is robust and effective for the classification and characterization of time intervals of changes in node and boundary maps. Our results reveal that changes in connectivity at the boundaries are slower and more complex than in those observed in the nodes, but of similar magnitude according to accuracy confidence intervals. These results were strongest in the posterior cingulate cortex and precuneus. As network boundaries are indeed under-investigated in comparison to nodes in connectomics research, our results highlight their contribution to functional adjustments to rTMS.

## 1. Introduction

Repetitive transcranial magnetic stimulation (rTMS) has become a popular method for the non-invasive modulation of brain function [1]. Recent neuroimaging studies have shown that functional changes induced by rTMS in a localized cortical region lead to selective and distinct modulation of activity and functional connectivity both within and between large-scale brain networks [2]–[7]. The mechanisms by which rTMS induces network modulations are still not well understood. Today, mapping whole-brain effects caused by local neural perturbations, including by rTMS, is a growing field of research. Well-established methods now allow for the assessment of connectome-level functional adjustments to high frequency rTMS in both node and boundary maps in sequential time intervals [8], [9].

Functional magnetic resonance imaging (fMRI) data obtained while participants are not engaged in any specific task is called resting state fMRI (rsfMRI). RsfMRI has been instrumental for advancing our understanding of the brain’s macroscopic functional network architecture [10]–[12] as well as which regions might be most functionally altered in psychiatric disorders [13], [14]. However, fMRI data typically consists of functional time-courses in thousands of voxels, which on the one hand allows for accurate inference of correlations or “functional connectivities” between regions, but on the other hand has high dimensionality. Different approaches have been proposed to reduce data dimensionality of the data and to identify the most relevant patterns of spatiotemporal organization in fMRI data. This is the case for whole-brain functional regions that will be represented in our study as nodes [15] and boundaries [8], [16], [17]. Nodes are defined as the greatest strength of local or global connectivity, also known as concepts of modularity and integration respectively, which enabled many insights into dimensional organization of the healthy and diseased brain [18]. Boundaries are the counterparts of the nodes, identified where the connectivity strength is the lowest or absent, usually in the transition between neighboring functional regions [16]. In contrast to the investigation of boundaries, the scientific community has given disproportionate attention to the nodes of functional networks. In node clustering approaches, spatiotemporal elements (i.e., voxels) may be grouped on the basis of the similarity versus dissimilarity of their functional connectivity [15]. An example of a node clustering approach is independent component analysis (ICA), which is used as a brain mapping method that efficiently segregates functional components based on their corresponding spatiotemporal distribution [19]. ICA has been widely applied to the identification of large-scale brain networks [5], [20], [21].

Group-based parcellation schemes use fMRI data from multiple individuals to map large-scale functional brain networks, that is, collections of widespread regions showing functional connectivity [17], [22]–[24]. While group-based parcellation captures major features of functional brain organization that are evident across individuals, such approaches may obscure certain person-specific features of brain organization. In contrast, subject-specific parcellation methods have been shown to effectively map aspects of functional organization that differ for particular individuals (e.g. Kong et al., 2019; Saxe et al., 2006). Several recent studies have demonstrated that extensive rsfMRI data collected across multiple sessions from the same individual can be used to delineate high-quality cortical parcellations at the individual level [26]–[28]. Subject-specific parcellation may enable increasingly precise planning and delivery of rTMS interventions.

Machine learning includes the development of algorithms that can detect spatially complex and often subtle patterns in highly dimensional data, and has been applied to neuroimaging. Machine learning may thus be useful for individual-level predictions that could ultimately be used in clinical contexts [29]–[32]. The support vector machine (SVM) is a machine learning algorithm that constructs hyperplanes in multidimensional space to optimally separate data classes [33]. Taken together, in the current study SVM was used to identify the strongest rTMS-related changes in brain functional connectivity patterns in both nodes and boundary maps. SVM is one of the most commonly used machine learning algorithms due to its robustness and easy interpretability. A particular advantage of SVM in this study is that it often yields better classification in smaller datasets, that is, datasets in which the number of features greatly exceeds the number of training data samples [34]. As a commonly applied resource, the classification task can be simplified by using unbiased feature selection approaches. Features selection involves the identification of the most useful data features in the training dataset [35], which are then solely used for classification. Features selection has been shown to improve accuracy while also increasing the interpretability of identified multivariate models [36], [37].

Recent neuroimaging and predictive modeling findings suggest that a locally generated brain stimulation-induced perturbation in neural activity is gradually integrated by selective alterations of within- and between-network dynamics [2], [38]. In addition, animal studies have shown that 10 Hz rTMS creates a transient cortical functional state that is characterized by increased excitability and increased response variability [39], [40] 10◻Hz rTMS applied to cat visual cortex resulted in a reduction of the inhibitory notch commonly seen in visual evoked activity, evidencing decreased inhibition during visual processing [40]. On the other hand, the findings by [39] implicate a reduction of specificity (decorrelation) close to the borders of the functionally distinct regions reflected in the widening of boundaries between them. This is plausibly happening due to rTMS-induced reduction of inhibition, predominantly in the boundaries but also in the nodes.

In the current project, we endeavored to understand rTMS modulation of whole-brain connectivity patterns. For that, we used individual-level comparisons to create sham-corrected maps for nodes and boundaries. This approach enhanced the sensitivity of detecting individual-level rTMS-induced variations in functional connectivity. Our SVM approach allowed for the data-driven identification of the most substantial functional changes as a whole and pairwise across time conditions (time intervals R). Based on the prior studies described above, we hypothesized that 10 Hz rTMS would affect functional connectivity both in the nodes and in the boundaries of distant regions that interact with the DLPFC.

## 2. Material and methods

### 2.1 Participants and study design

23 healthy subjects between the ages of 18-65 were recruited from a university environment to participate in a double-blind, sham-controlled, crossover design study that investigated the neural effects of 10 Hz rTMS using rsfMRI. Further details on the study design have been reported elsewhere [41]. Participants were screened with a self-report clinical questionnaire and the Symptom Checklist 90 Revised (SCL-90-R) to ensure that they had no current or previous history of neurological or psychiatric disorders. Additional exclusion criteria included recreational drug use in the past month, current or history of substance abuse or addiction, any contraindications to MRI or TMS (e.g., pregnancy, epilepsy), history of traumatic brain injury, participation in any TMS or ECT study in the past 8 weeks, and unwillingness to consent or to be informed of incidental findings. Informed consent was obtained from all subjects before their inclusion in the study. The study protocol was approved by the Ethics Committee of the University Medical Center Göttingen (UMG). This study was conducted in accordance with the current version of the Declaration of Helsinki.

For each participant, experiments were conducted over the course of 3 visits with approximately one week in between each visit. As described in [41], on visit 1 a structural T1-weighted volume and rsfMRI were acquired for the identification of a subject-specific DLPFC site for rTMS stimulation. This target was then used for real and sham stimulation protocols on visit 2 and 3. At the beginning of visit 2 and 3, an rsfMRI scan (R0) was obtained pre-rTMS. Next the resting motor threshold (RMT) for each subject and session was determined, which was then used to set the stimulation intensity (i.e., 110% of the RMT). Thereafter, a 10 Hz rTMS clinical protocol of 3000 pulses was delivered over 37.5 min. This procedure was additionally controlled by sham rTMS in a double-blind counterbalanced crossover design. RTMS was precisely delivered to each subject at the pre-selected DLFPC target, guided by online neuronavigation. Three additional rsfMRI scans were obtained post-rTMS at 10-15 min (R1), 27-32 min (R2), and 45-50 min (R3) after the end of stimulation (non-continuous time slots of about 5 min each). These acquisitions allowed for the assessment of functional connectivity changes induced by rTMS.

### 2.2 Neuroimaging data acquisition and preprocessing

Structural T1-weighted scans with 1-mm isotropic resolution and functional data were obtained using a 32-channel head coil and 3T MRI scanner (Magnetom TRIO, Siemens Healthcare, Erlangen, Germany). For rsfMRI, 125 volumes were acquired in approximately 5.5 minutes using a gradient EPI sequence with the following parameters: TR of 2.5 s, TE of 33 ms, 60 slices with a multiband factor of 3, FOV of 210 mm × 210 mm, 2×2×2 mm, with 10% gap between slices and anterior to posterior phase encoding.

RsfMRI preprocessing was conducted in Data Processing Assistant for Resting-State fMRI software (DPARSF V4.4, [42]). Initial steps of preprocessing included the slice timing and head motion correction [43], [44]. Afterwards corrected images were analyzed using the SPM12 gradient echo field map unwarping tool [45]. White matter, CSF, and global signal were then regressed out to additionally reduce nuisance effects [46]. Corresponding T1-weighted images were adjusted to the standard Montreal Neurological Institute (MNI) template and then used for spatial normalization of rsfMRI with a resampled voxel size of 3×3×3 mm. The preprocessed images were then spatially smoothed with a 4×4×4 mm full width at half maximum (FWHM) Gaussian kernel. Data were then detrended and band-pass (0.01–0.1 Hz) filtered to remove bias from low-frequency drift and high frequency noise.

### 2.3 Parcellation of individual-level resting state functional correlations

Subject-related parcellations were obtained using two methods: (1) the “Snowballing” algorithm provided a connectivity node density map [9] and (2) Boundary Mapping generated an average spatial boundary map [8], [9]. Both methods use whole-brain resting state functional connectivity (RSFC) to create individualized three-dimensional node and boundary maps. Briefly, the Snowballing algorithm uses seed-based RSFC to identify locations that are correlated (the “neighbors”) with a starting seed region of interest (ROI) location. These neighbors are then used as the new seed regions and the procedure is repeated iteratively. Each iteration of identified neighbors in this procedure is referred to as a “zone”. The neighbors correlated with a given seed region are spatially clustered with their distinct local maxima. Averaged peak voxel locations over multiple zones results in a node density map that represents the number of times a voxel was identified as a node across all ROI correlation zones from starting seed locations. As in [9], nodes were identified as the peak distribution from three zones of Snowballing. In the original study the snowballing parcellation was performed on a cortical surface. Here, we extended it to the whole-brain volume space to include subcortical regions and to match the dimensionality of our feature selection methods. As this has not been shown before, we have validated 3D node density maps by showing that inter-individual variability exceeds intra-individual variability both in the current dataset as well as in an independent dataset (**Supplementary Figure 2, left panel**). Most importantly, these node distribution maps have been shown to be invariant to the locations of starting seeds as well as the seed sizes [9]. While in the original study 264 seeds were used, our 278 initiation points corresponded to 278 ROIs of 5mm radius, containing additional subcortical and subcallosal seeds [10]. The correlation maps were thresholded at r>0.3.

In contrast to the definition of nodes, RSFC-Boundary Mapping identifies locations where the patterns of RSFC correlations exhibit abrupt transitions, and therefore provides estimates of putative boundaries between functional regions [8]. In contrast to the initially proposed technique that flattens a given subject’s cortex into a 2D surface [9], we performed all computations directly on the subject’s full-volume 3D Cartesian grid. This allowed us to simultaneously overlay, for each subject, the 3D Snowballing node density images with the RSFC-Boundary Mapping output (**Figure 1**). Similar to full-volume nodes, full-volume boundaries were validated by showing that inter-individual variability surpassed intra-individual variability (**Supplementary Figure 2, right panel**). In line with the original report, we also found that boundaries resulting from RSFC-Boundary Mapping (**Figure 1, regions in green**) typically surrounded the nodes of high peak count identified by the Snowballing algorithm (**Figure 1, regions in red**). Aside from this difference, we followed the RSFC-Boundary Mapping method as previously described [9]. To speed-up computation time, the full-resolution preprocessed fMRI data were first resampled to a coarser grid, resulting in 19,973 voxels (4.5×4.5×4.5 mm in size). The whole-brain RSFC map was then computed for each voxel by correlating the time series of the given voxel with every other voxel. Then, the similarity between each voxel’s temporal correlation map and every other voxel’s temporal correlation map was computed as the spatial correlation between them. That resulted in 19,973×19,973 symmetrical spatial correlation matrices. Every row of this matrix corresponds to a reference voxel and can be displayed in the volume of the brain where the value in every voxel is the similarity between the temporal correlation map at that position and the reference voxel. Every row was then mapped to the brain mask back, representing voxel’s similarity map. Next the spatial gradient was computed and then averaged across all similarity maps. The average of those spatial gradient maps represents the probability with which each location is identified as a point of rapid change in the RSFC maps between two neighboring areas.

**Figure 1:**
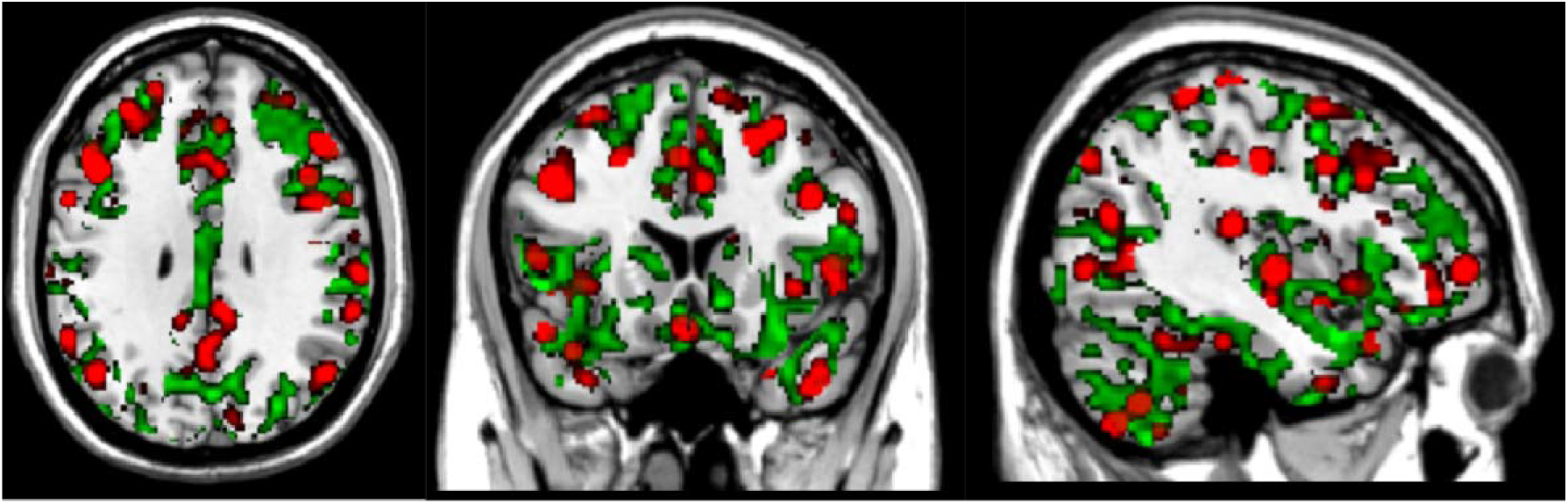
An example of individual-level node and boundary maps. It is noteworthy that calculated RSFC-Snowballing nodes (red areas, thresholded r=0.3) are often surrounded by transitional areas represented by RSFC-Boundary mapping boundaries (green areas thresholded r=0.08), revealing the complementary nature of both parcellation methods.

### 2.4 Feature selection

Machine learning models can be directly applied to both node and boundary maps. However, due to a large number of features (104-105 voxels) compared to the number of subjects in this study (n=23), that would lead to a substantial overfitting. Subsequently, overfitted classification models suffer from unsatisfactory interpretability and accuracy [47], [48]. In order to select a smaller number of physiologically plausible voxels located in node and boundary maps, we employed a novel data-driven feature selection approach based on the resting state networks (RSNs). RSNs were accessed in the same group of subjects but from an independent dataset (Visit 1) from the classification dataset (Visit 2 and 3). The RSNs were identified by group ICA [11], [47] using MELODIC tool of FSL 5.0.7. We temporally concatenated the fMRI data of 23 subjects recorded on visit1. Based on visual inspection and the power spectrum of the MELODIC output, we selected the nine best-fitting spatiotemporal independent components (IC). While the node regions were assumed to lie close to maxima of the selected ICs, the boundary areas were rather identified at the intersections of the ICs.

As the different ICs showed different strengths (distributions of values across the brain) and we intended to have multiple networks represented rather than a single “winner”, the node areas were defined by overlaying all ICs. The percentage of voxels belonging to every IC was controlled by the threshold; voxels with lower strength were discarded. Next, the components were merged together resulting in an array of nodal voxels. The value of the node density (Snowballing) map of each subject/time interval was then extracted for each of those voxels. This process was designed to select the most important features that would then be modeled using the classification algorithm. An example set of node masks based on all subjects, but independent from the dataset used for classification, is presented in **Figure 2**, left panel (thresholded at 97.5% and 99% of discarded voxels).

**Figure 2:**
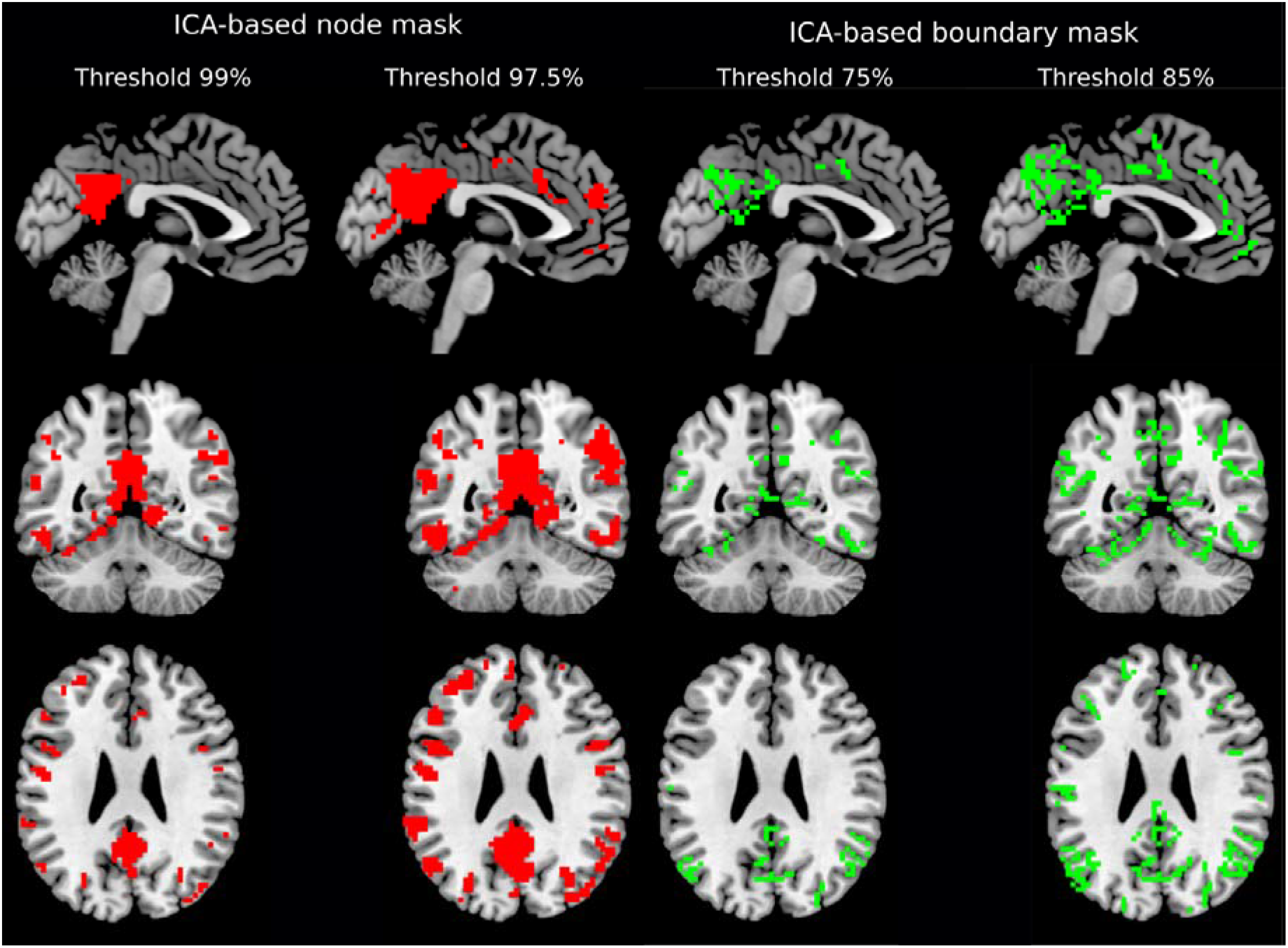
ICA-based binary masks of node (left) and boundary masks (right) from a dataset not used in machine learning classification. ICs node regions grow in size when one increases the threshold value. At the moment two or more ICs meet in one particular voxel, that voxel is identified as the boundary. Changing the threshold value allows for the optimization of the number of voxels corresponding to both masks.

In case of the boundary mask, we have developed an algorithm closely related to watershed transform [49]. The underlying idea of watershed transform is finding an optimal position for dams to be built, where the water coming from different basins, according to the surface shape, would meet. In our case, the surface is represented by all ICs and the basins are the strongest voxels according to ICs. Starting from the maximum of each component (100% threshold), and then reducing the threshold in 1% steps, the ICs increase in size. A voxel is identified as a boundary when two or more ICs intersect each other (i.e., the same voxel is included in multiple ICs). Single voxels, i.e. not surrounded by more voxels from the same IC, were not treated as boundaries to avoid spurious findings. The number of included voxels were dependent on the percentage level step that was used to descend from the maximum. An example set of boundary masks based on all subjects, but independent from the dataset used for classification, is presented in **Figure 2**, right panel (threshold values of 75% and 85%).

### 2.4 SVM classification

The capacity of SVM to predict outcomes of the rTMS intervention was first assessed using leave-one-group-out stratified cross-validation (CV), where the group was defined as all of stimulation conditions (time intervals R) of the same subject. Our main analysis employed SVM for multiclass classification with “one-versus-one” approach of all sham-subtracted conditions, i.e. R0 vs R1 vs R2 vs R3. SVM classification was performed using a linear kernel and the default scaling factor of 1. Sex and age were not regressed out because in every comparison both groups were represented by the same subjects, and thus automatically balanced.

The number of features (voxels) used as an input for SVM varied by changing the threshold value of the extracted ICA-based node and gradient masks (**Figure 2**). SVM assigns weight to every voxel, which can be interpreted as an importance of the voxel to separate conditions. The SVM was trained for every pair of conditions starting with a mask threshold corresponding to 10,000 voxels. By changing the threshold for both ICA-based node and gradient masks, we removed the “weakest” voxels and trained the SVM again from scratch. This procedure was repeated until the total number of voxels surviving thresholding reached zero. This recursive process allowed for the assessment of a model accuracy curve that was defined by the percentage of included voxels. To access the confidence interval of the accuracy curve, we ran SVM on 1000 bootstrap samples. The global maximum of this model mean across bootstraps accuracy curve represents the most informative set of voxels for classifying all of the conditions. In case the accuracy curve (or a certain part of it) is consistently higher than the chance level (>25%), we perform pairwise classification, i.e. R0 vs R1, R0 vs R2, R0 vs R3, R1 vs R2, R1vs R3 and R2 vs R3, to identify the time and the direction of the most significant changes happening after 10Hz rTMS. The same algorithm was applied to both the node density and the gradient maps across voxel thresholds. This procedure is schematized in **Figure 3**.

**Figure 3:**
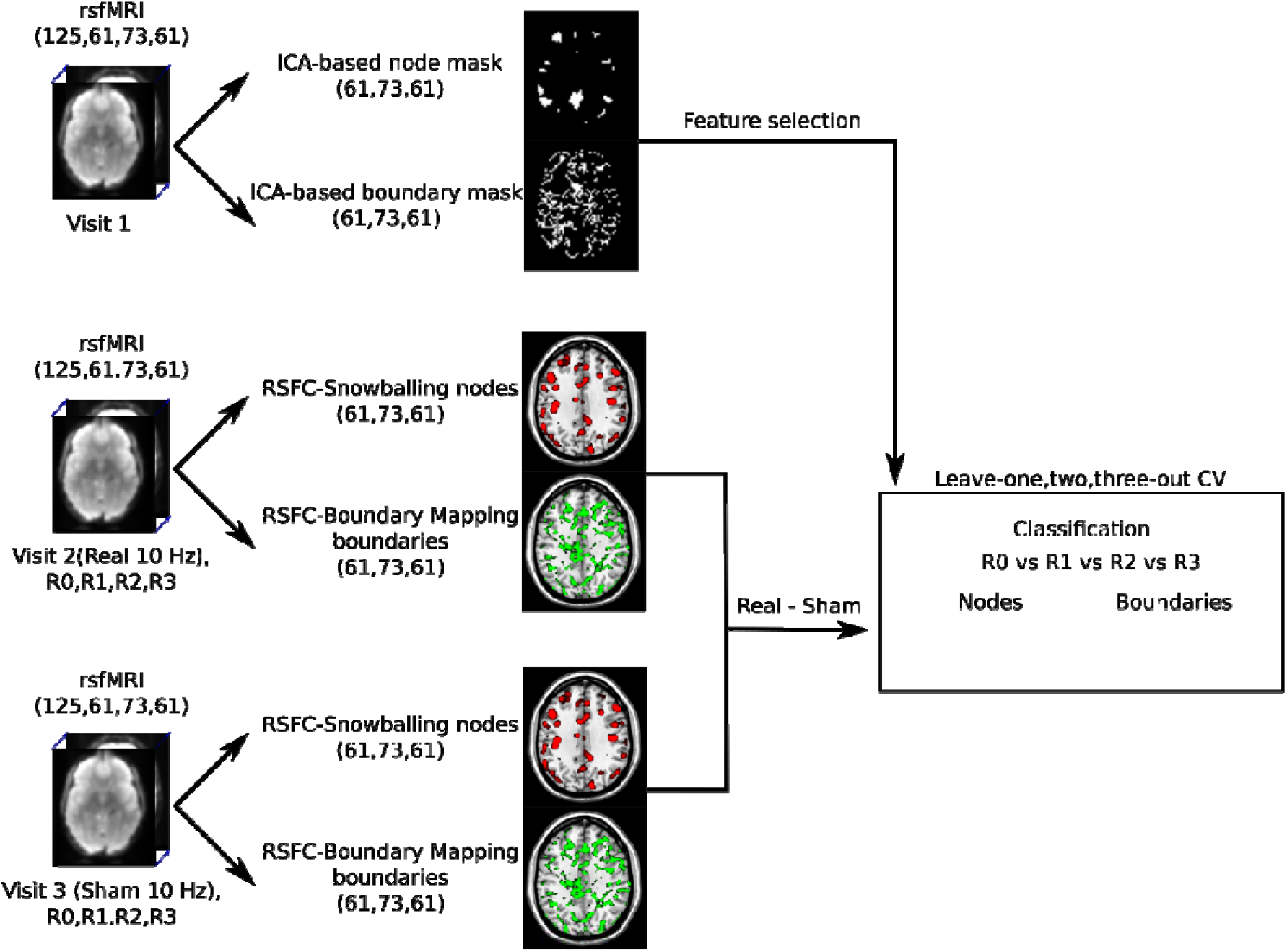
Schematic diagram of the analysis pipeline. RsfMRI data from 4 sessions (R0, R1, R2, R3). Both rsfMRI from Visit 2 (real rTMS) and Visit 3 (sham rTMS) were used to compute RSFC-Snowballing density nodes and RSFC-Boundary Mapping gradients. Next sham stimulation maps were subtracted from the corresponding real stimulation maps. The extracted ICA-based masks derived from the visit1 measurements were then applied to the corresponding node and gradient maps for feature selection. Finally, the remaining voxels were used for machine learning classification.

CV is well known as an effective method in machine learning for testing generalizability of a trained model. Model performance was further tested using leave-two and leave-three-out stratified CV (7 fold and 11 fold CV) in which 2 or 3 subjects were withheld from training and assigned to the test sample. Our goal with this approach was to investigate the effect of the amount of data provided to a classifier, and thereby assess the impact of CV strategy. By providing more training data, as in leave-one-out CV, the model has more generalizable performance compared to leave-two-out and leave-three-out CV [50]. The computations were completed in Python using custom-written scripts that used functions from the Nilearn v0.7.0 (https://nilearn.github.io/) and Sklearn v0.23.2 (https://scikit-learn.org/stable/) libraries.

### 2.5 Effect of head motion on the separation of conditions

To exclude the possibility that condition-related differences are caused by head motion [51], we performed pairwise classification of mean frame-to-frame head displacement across real, sham, and real-sham conditions (R0, R1, R2, R3). The resulting SVM performance accuracies are presented in **Supplementary Table 2**. As the performance of algorithms based on head motion are close to chance, this analysis confirmed irrelevant influences of head motion on the results of classification based on RSFC nodes and boundary maps. Additionally, a cutoff to remove the subjects with high-motion frames according to the threshold (mean FD = 0.5) was set, yet no session surpassed this value.

## 3. Results

### 3.1 RSFC nodes’ density maps

We performed SVM classification on RSFC nodes density maps across all conditions and applied an iterative ICA-based feature selection step to spatially identify nodes that were strongly modulated by 10 Hz rTMS. The threshold for the ICA-based node mask, corresponding to the percentage of voxels removed, varied between 99.9% (154 voxels) to 94.5% (10,799 voxels) in 0.1% increments. The highest accuracy of 33.2±0.8% was achieved for a threshold of 99.1%, yielding 1690 voxels fed to the SVM (**Figure 4a**). Results indicated that by increasing the threshold, many informative voxels for the classification were discarded, yielding the massive accuracy drop between 99.9% and 99.7%. Lower thresholds led to a decrease in accuracy, which may be due to model overfitting (i.e., inclusion of uninformative voxels). Additionally, we compared the performance of the SVM model with a different number of CV folds. All three CV schemes showed similar accuracies across thresholds ranging from 99.9% to 98.7%. Lower thresholds for leave-three-out CV resulted in slightly higher accuracies compared to leave-one-out and leave-two-out CV. The most informative voxels were located in the posterior cingulate cortex, angular gyrus, anterior insula, and visual cortex (**Figure 5, top panel**).

**Figure 4:**
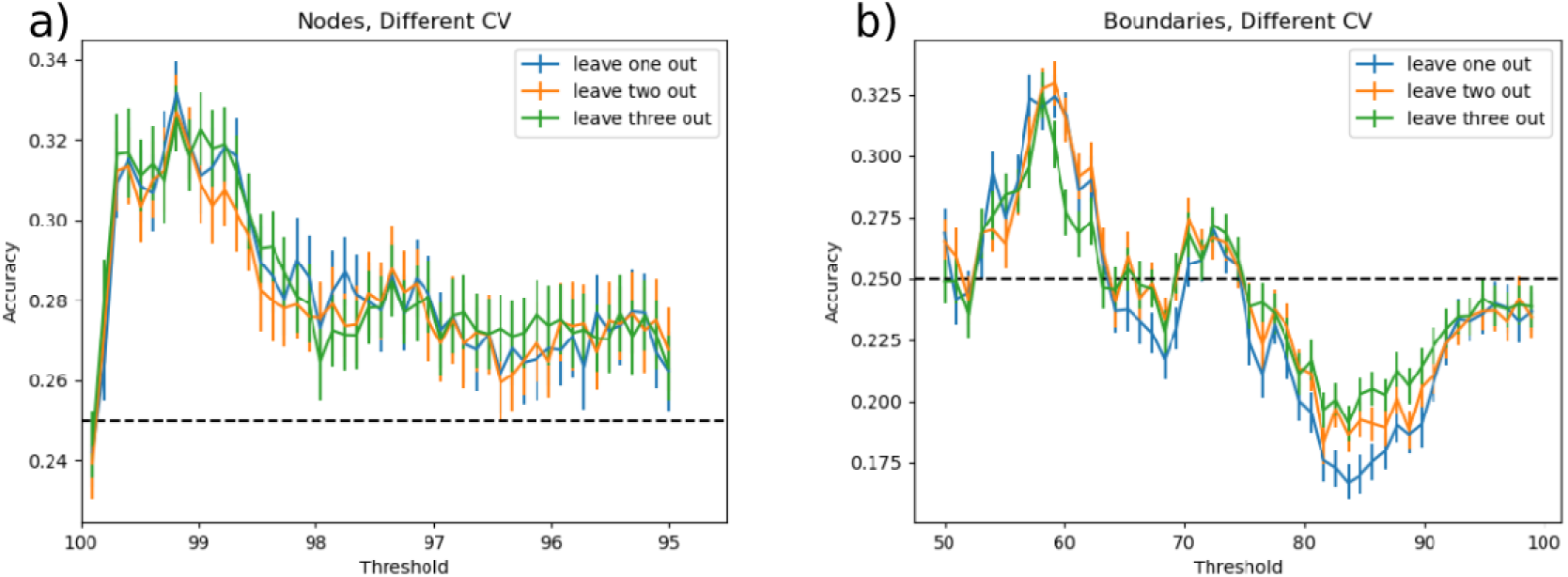
Multiclass classification accuracy of SVM using node density map (**a**) and boundaries (**b**) across all threshold levels. The error bar indicates 95% confidence intervals over 1000 bootstraps. To test the stability of classification results, classification was also performed for leave-one-, -two- and -three-out cross-validation (CV) routines. The highest performance for peaks (Acc =33.2%) was achieved for the threshold value of 99.1% corresponding to 1690 informative voxels. For boundaries, the threshold of 59% yielded the highest performance (Acc=32.4%) resulting in 113 voxels to be strongly modulated via 10Hz rTMS. Dashed line represents random-choice accuracy, one of four R categories.

**Figure 5:**
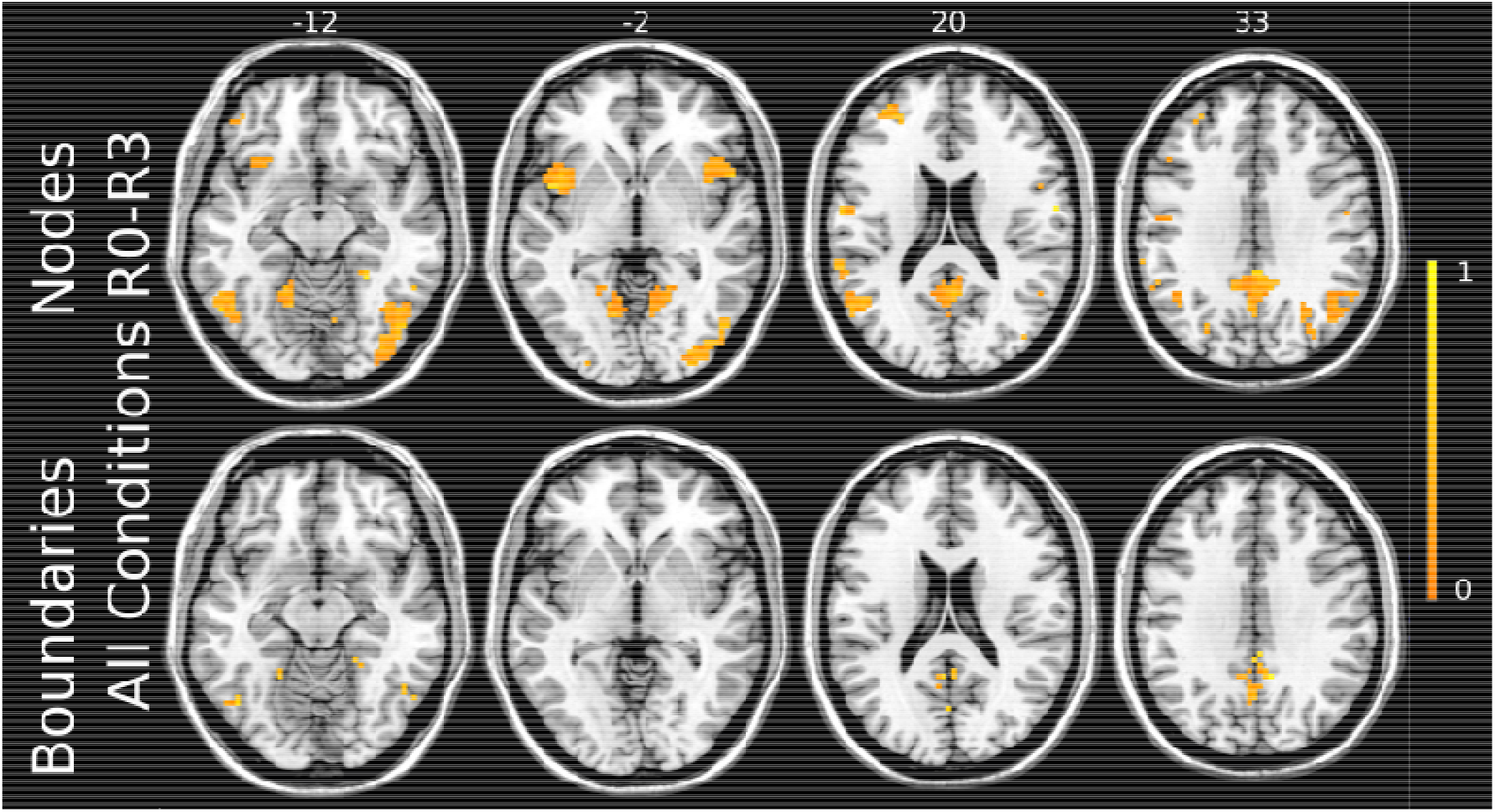
Most strongly modulated voxels corresponding to the highest multiclass classification accuracy of SVM for both nodes and boundaries. The strength of modulation is color-coded by a warm colormap over the individual template. The top numbers refer to axial plane z-coordinates in MNI space.

Since the accuracy was consistently higher than by chance, we aimed to discriminate the sessions in which connectivity was most strongly modulated by 10 Hz rTMS by performing pairwise classification of conditions. The highest accuracy of 74.2±1.1% was achieved for R1 vs R2 comparison with a threshold value of 99.7%, yielding 764 voxels used in SVM-based classification (**Figure 6a**). All three CV schemes yielded similar accuracies for R1 vs R2 classification across thresholds ranging from 99.9% to 98.7% (**Supplementary Figure 3**). Majorly, the same set of voxels with both positive and negative weights was found modulated in R1 vs R2 classification (**Figure 7, top panel**) as in multiclass classification.

**Figure 6:**
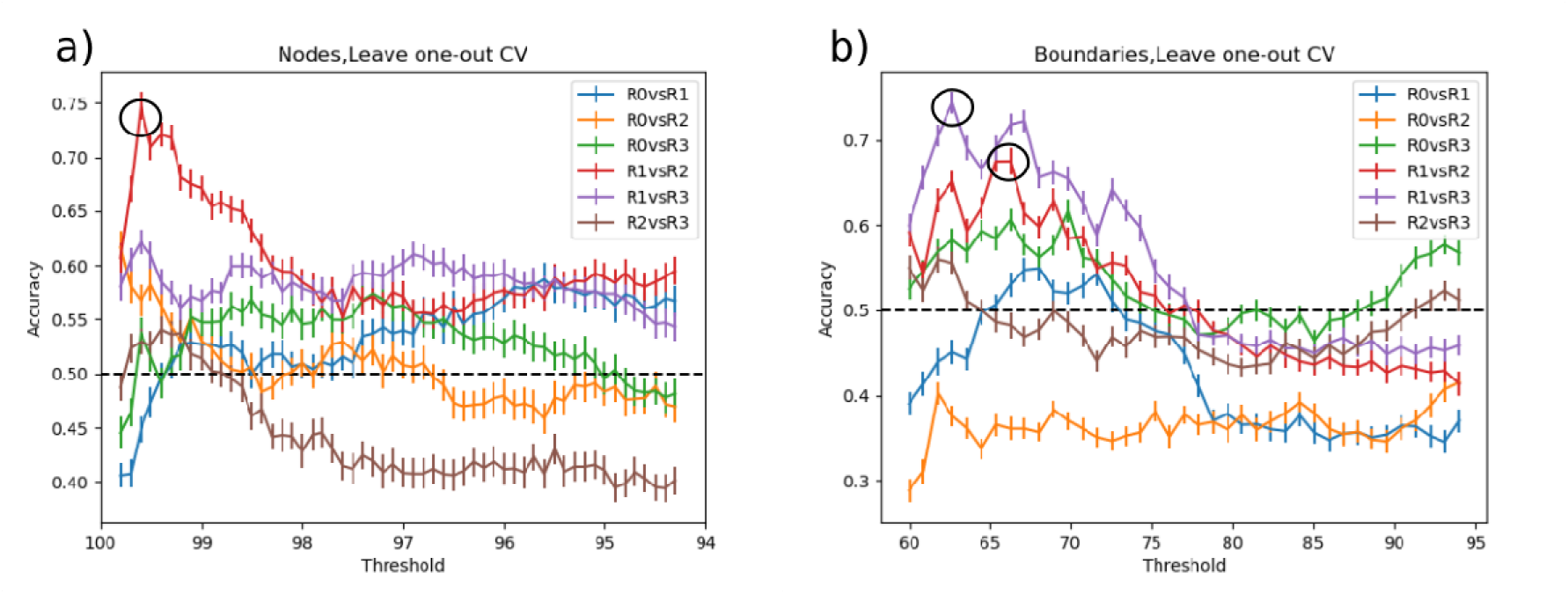
Pairwise classification accuracy of SVM using nodes (a) and boundaries (b) across all threshold levels. The highest accuracy for nodes was achieved by R1 vs. R2 comparison for the threshold value of 99.7%, corresponding to 764 voxels. In case of boundaries, R1 vs. R3 yielded the highest accuracy for the threshold value of 63% (174 voxels). The second highest accuracy was obtained by R1 vs. R2 comparison for the threshold value of 67% resulted in 346 voxels being highly modulated by 10 Hz rTMS. Dashed line represents random-choice accuracy, one of two R categories.

**Figure 7:**
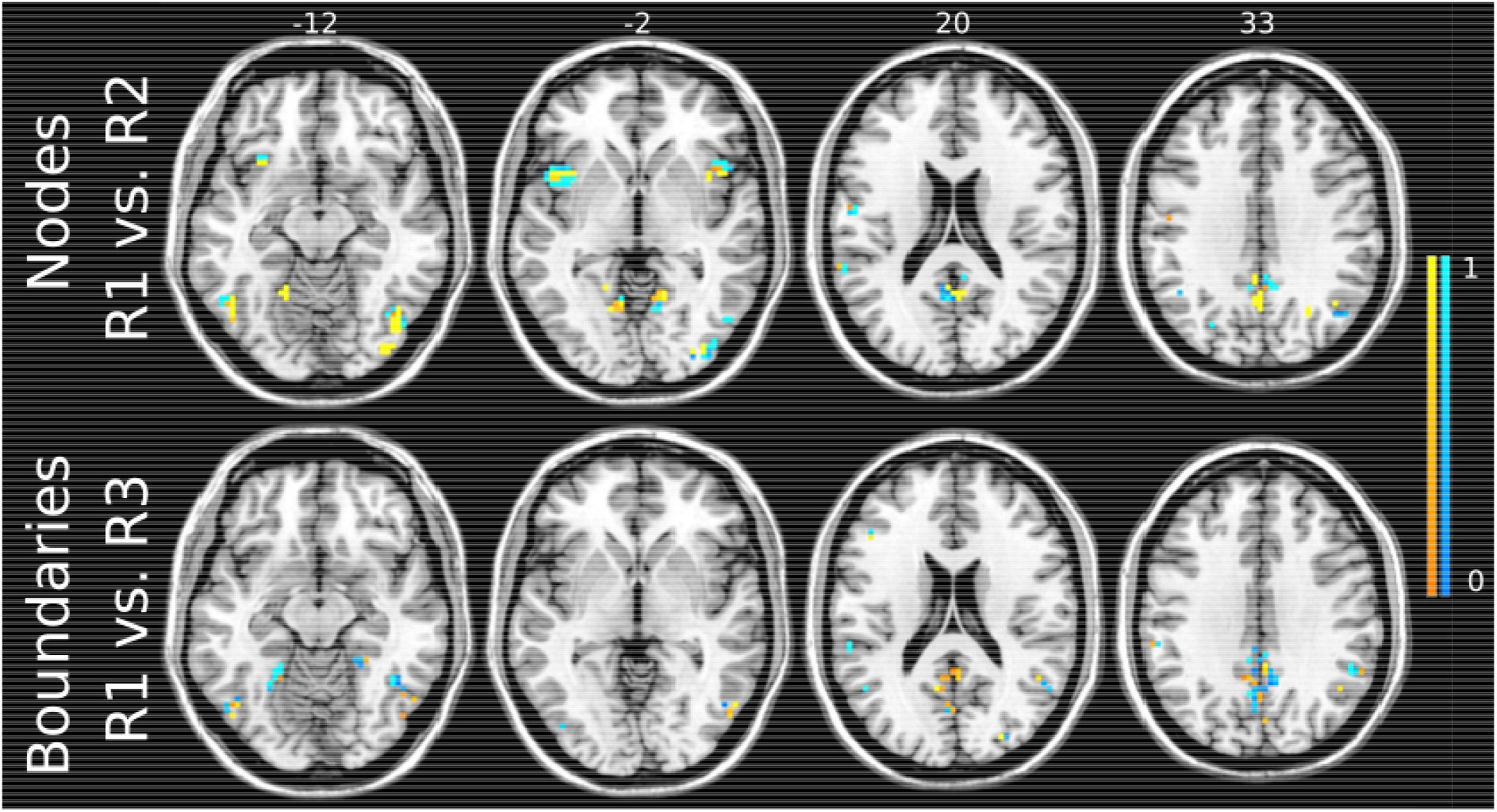
Voxels corresponding to the highest performance of the SVM using the Snowballing nodes for R1 vs R2 comparison (top) and the Boundary-Mapping boundaries for R1 vs R3 comparison (bottom). The sign and strength of modulation is color-coded by warm colormap (connectivity increase) and cold colormap (connectivity decrease). The top numbers refer to axial plane z-coordinates in MNI space.

### 3.2 RSFC boundary maps

For boundaries extracted from RSFC-Boundary Mapping, the threshold for ICA-based boundary masks was varied from 50% (23 voxels) to 94% (10861 voxels) in 1% increments.

The highest accuracy of 32.4±0.9% was achieved by the threshold of 59% corresponding to 113 voxels being fed into the SVM (**Figure 4b**). These voxels were predominantly located in the ventral posterior cingulate cortex, precuneus, angular gyrus, and fusiform gyrus (**Figure 5, bottom panel**). Consistent with the analysis on Snowballing nodes described above, increasing the number of voxels extracted from the ICA-based boundary mask caused a drop in the overall accuracy of the SVM. All three CV routines showed a similar pattern in accuracy across the threshold range of 50%-67%.

Similar to the analysis based on RSFC nodes’ density maps, we performed pairwise classification of conditions to detect the timing of the strongest changes in boundaries. The highest SVM accuracy of 74.5±1.1% was achieved for R1 vs R3 (**Figure 6b**) for the threshold value of 63% (174 voxels, **Figure 7**), Using leave-one-out CV resulted in the performance improvement from 72.3% to 74.5% compared to leave-two-out CV for the ICA-based boundary mask threshold of 63% (**Supplementary Figure 4**). The learning curve of R1 vs R3 across CV approaches (**Supplementary Figure 4**) indicates the highest masking threshold at which leave-one-out CV strategy yields the highest (or at least equal accuracy compared to leave-two and leave-three-out CV) was 76%. This masking threshold corresponded to 1194 voxels. For higher threshold values, leave-two-out CV exhibited slightly higher accuracy than the other approaches.

The second highest accuracy of 68.5±1.3% was achieved by R1 vs. R2 classification (**Figure 6b**) using a threshold value of 67% that corresponds to 346 included voxels. For threshold values from 60% to 86%, both leave-one-out and leave-two-out CV strategies were associated with higher accuracy compared to leave-three-out CV (**Supplementary Figure 4**). The sign of feature weights in the majority of identified voxels was in the opposite direction to the R1 vs R2 and R1 vs R3 comparison (**Figure 8a and b**). The highest classification results are presented in **Supplementary Table 1**.

**Figure 8:**
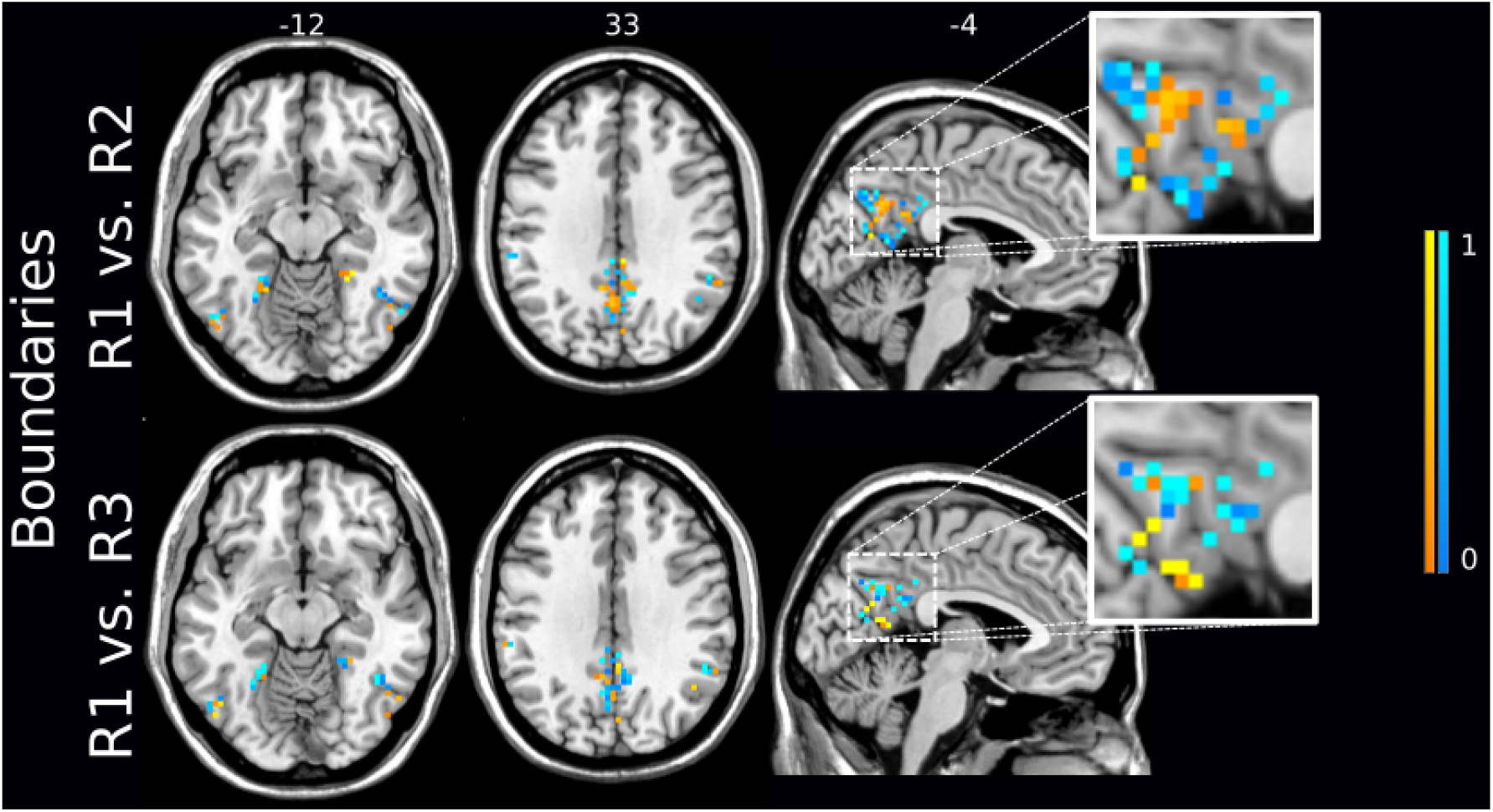
Two highest classification accuracies based on the Boundary-Mapping boundaries were achieved for comparisons R1 vs R2 (top) and R1 vs R3 (bottom) with 329 and 174 stable voxels respectively. The sign and strength of modulation is color-coded by a by warm colormap (connectivity increase) and cold colormap (connectivity decrease). The zoomed-in image shows that the majority of discriminative voxels in both comparisons were located in the posterior cingulate cortex (PCC) and precuneus. The top numbers refer to axial plane z-coordinates in MNI space.

## 4. Discussion

There is considerable interest in understanding the changes in functional connectivity driven by 10 Hz rTMS, both in the field of neuroscience and in applied clinical practice. Here we have used a predictive SVM model approach to identify the locations and time intervals after stimulation that exhibit the most substantial whole-brain functional changes in large-scale network nodes and boundaries. To overcome the issue of high-dimensional data, that is, the number of voxels in node and boundary maps being much greater than the number of subjects in the study, we have applied feature selection using multiple threshold ICA-based masks created from separate fMRI data. Confirming our hypothesis, we have identified connectivity changes related to 10 Hz rTMS in both network node and boundary maps. Of note, as indicated by cross-validated SVM, changes majorly comprise the PCC, angular gyrus, insula and fusiform gyrus. Accuracy confidence intervals in the classification of boundaries are similarly as substantial as those occurring in nodes. A complex pattern of changes was observed in boundaries alternating between decreases and increases in functional connectivity, which was particularly evident but not limited to the PCC and precuneus.

### 4.1 Individual nodes and boundaries

This work extends to the concept of modularity and integration frequently addressed in the field of connectomics [18],, both of which focus on the perspective of nodes in the formation and reorganization of networks in cognitive functions and in clinical disorders. Most connectomics studies to date have been focused on the strongest points of functional connectivity, “hubs”, and the way that such hubs are organized to efficiently propagate information across regions [52], [53]. Transition gradients have recently received more attention in the literature [54]–[56]. To the best of our knowledge, the current study is the first to show which changes occur in both functional connectivity nodes and boundaries after 10 Hz rTMS. Indeed, based on their accuracy and confidence intervals, we see that rTMS effects related to boundaries are similarly as substantial as those seen in nodes. Cytoarchitectural divisions of the PCC into dorsal and ventral parts have been previously shown to exhibit distinct functional connectivity patterns [57]. According to findings from a recent study [58], both PCC and precuneus contain predominantly transient nodes - an entropy class that the authors used to classify changes in network assignments across subjects and brain states. Our findings are consistent with this notion, highlighting that further functional heterogeneity is largely elicited in these regions, particularly in boundaries, after 10 Hz rTMS.

The exact function of boundaries is not yet well understood, which might have discouraged its systematic evaluation. It may be speculated that boundaries may act to segregate information within functional regions, or that they may support network stability [59]. Another possible role of boundaries may be supporting functional adaptation of a given region during plastic changes in the mature primate brain. Evidence for this has been shown in primary motor and sensory regions (for review see Florence et al., 1997). In support of this theory are several studies that have shown that rTMS-induced changes in neuronal inhibition can prime cortical networks for the expression of subsequent experience-dependent plasticity [39], [61], [62]. The high frequency rTMS potentially creates a cortical state with enhanced plasticity, opening a time window for targeted re-learning of connectivity patterns [39]. A complex pattern of changes in the functional connectivity of boundaries, which was particularly evident but not limited to the PCC and precuneus, is reminiscent of the neuronal population dynamics in the cat visual cortex after 10 Hz rTMS perturbations [40]. In the animal experiments, stimulation induced initial suppression of on-going cortical activity, followed by an increase in cortical excitability that lasted about 2-3 hours, but was prone to slow activity fluctuations. Worth mentioning that this animal study as well as clinical TMS/EEG findings reported by [63] suggest a different mechanism than excitatory LTP. Indeed, rTMS may instead reduce the local intracortical inhibition leading to long-lasting neuromodulatory effects in both the boundaries and the nodes.

Causal effects of 1 and 5 Hz rTMS on global functional connectivity have been explored recently using fMRI [64], [65]. These studies have found that distant effects, that is, effects relatively far from DLPFC, are determined by connectivity profiles of the stimulation target with macroscopic networks. Excitatory 10 Hz rTMS in the left DLPFC, as we used in the current study, resulted in multivariate patterns of increases and decreases in functional connectivity. These fluctuations occurred primarily in the PCC, angular gyrus, fusiform gyrus, and insula regions. Each of these regions have been previously shown to be functionally related to the DLPFC [66]–[69], reinforcing the notion of distant effects of this stimulation protocol. In the current study, the SVM most substantially identified functional connectivity changes in nodes that occur about 30 min after stimulation, in line with a previous study performed on the same dataset, but using factorial design ANOVA to find group differences [41]. In contrast, effects related to boundaries were temporally extended up to 45 min. Previous studies that investigated the effects of 10 Hz rTMS with rsfMRI constrained their analyses to particular seeds or networks of interest. Another important advancement provided by the current study is an assessment that started from a global evaluation of 10 Hz rTMS effects, considering maps of nodes and boundaries at the subject level at different intervals. The importance of individualized characterization of functional brain networks has been highlighted in the literature [26]–[28]. These developments may inform future clinical applications based in “precision” or “systems” medicine. Toward this goal, methods applied in the current study might have been advantageous, considering that individual subject analysis of node and boundary maps closely correspond to the original study [9]. In our study though functional boundaries were not only extracted for the cortical surface. To match the resolution and dimensionality of the node maps, we calculated boundaries on the whole brain. Furthermore, our study is unique in that we used pairwise comparisons of each individualized map type, that is, nodes and boundaries, in the context of a double-blind design controlling for placebo effects. This may have allowed us to identify the contribution of boundaries to the functional changes caused by 10 Hz rTMS. This individualized approach was followed by SVM, which might have contributed to the identification of the most important effects related to the time intervals and areas.

### 4.2 SVM classification and feature selection

SVM is a machine learning approach with clear advantages over univariate models. With that said, important preconditions had to be fulfilled to avoid potential biases. One of the main concerns when we applied SVM was to prevent overfitting. The full-resolution rsfMRI was first resampled to 19,973 voxels to reduce the computation time of the spatial correlation matrix – the most computationally demanding step in the algorithm. Splitting the rsfMRI signal into two complementary maps also had the practical benefit of enabling independent assessment using SVM. This also reduced the number of input variables to the model, and thus further prevented overfitting. Secondly, the large number of voxels in both individualized node and boundary maps required additional feature selection. While there are several masking algorithms that can locate functional nodes of the brain, to our knowledge, there is no complementary boundary parcellation method associated with any of them. For this reason, we have used group ICs to create both nodal and boundary masks.

Voxels included in the masks are controlled by the threshold. While these masks were built in a complementary manner, these masks included common voxels in several regions, including ventral PCC, the boundary area between angular gyrus and fusiform gyrus, and the boundary area between fusiform gyrus and associative visual cortex. These regions are consistent with findings from a study that reported on low stability of connectivity regions [58]. Some of these regions have been associated with myelin content, particularly the ones that myelinate latest during development [70], [71]. As some of the implicated regions overlap with those with late myelination, this suggests a role for flexible connectivity in learning [65].

Our approach is closely related to the recursive feature elimination technique that has been previously applied in neuroimaging [50], [72]. In our method, voxels/features are not removed based on their rank according to SVM, but rather based on the information obtained from the independent dataset (not included in SVM modeling). Therefore, our approach is less prone to overfitting and inflating performance accuracy. On the other hand, a drawback of our approach is that some fraction of the voxels not included might be potentially informative for classification.

Considering our small sample size, the performance of the classifier was low – close to chance – when all the features were used. Since in the multiclass classification SVM has shown the accuracy consistently higher than by chance, we performed post-hoc pairwise classification of conditions to pinpoint the timing of the most significant changes. For most of the pairwise comparisons, there was a substantial drop in the accuracy when more than 1000 voxels were used in the model. This observation may be explained by the presence of too many insignificant voxels and a small sample size. After the optimal number of voxels was reached from unbiased feature selection, further removal of voxels resulted in a drop of the accuracy. This is consistent with previous studies that have used similar machine learning approaches [50], [73]. Moreover, we investigated estimated accuracy curves for different CV methods as the number of CV folds may influence the accuracy of machine learning. That is, threshold values that yielded higher accuracy for leave-three-out, in contrast to leave-one-out CV, are more likely to have inflated accuracy as the training set is smaller. In general, the trained model should underperform or at least yield similar accuracy when trained with less data, otherwise the performance of the model is ambiguous.

### 4.3 Limitations

Our work has important limitations that should be considered. As mentioned above, with feature selection not all information derived from both node and boundary maps was considered for classification. Therefore, we cannot exclude that additional areas might have been modulated by 10 Hz rTMS and, due to the feature selection procedure, are not identified in our results. While systematic evaluation of thresholds is a common approach in machine learning, we acknowledge that nodes and boundary masks behave differently when changing threshold values. This may have contributed to the exclusion of regions as part of boundaries at a faster rate than nodes. One possible explanation for this is non-uniform distance between networks, which may cause proximal areas in boundaries to interact/overlap at lower thresholds, compared to more distant regions in nodes. In addition, a larger sample size may have enabled further methodological improvements such as: (1) increasing the resolution in the masking procedure; (2) consideration of more voxels from the masks; (3) application of non-linear predictive algorithms; (4) better fine-tuning of classifier hyperparameters; and (5) an independent validation dataset to strengthen the reliability of our results.

## 5. Conclusion

Our findings provide evidence that SVM classifiers using ICA-based feature selection can identify different spatial definition, direction, and timing in the pattern of fMRI-based brain connectivity changes in functional nodes and boundaries derived from 10 Hz rTMS applied to the left DLPFC. Identified nodes and boundaries are located predominantly in the ventral PCC, precuneus, insula, and fusiform gyrus and appear approximately 30 minutes after the stimulation is performed. By dividing the signal into two complementary parts (nodes and boundaries), we highlight the contribution of boundaries to modulatory effects of high frequency rTMS. These findings provide new insights into the currently unknown role of boundaries in network organization, motivating future, related investigations for the advancement of connectomics.

## Data Availability Statement

The algorithms and codes that were used in this study are available from the corresponding author upon reasonable request.

## Acknowledgements

Funding: This work was supported by the German Federal Ministry of Education and Research (Bundesministerium fuer Bildung und Forschung, BMBF: 01 ZX 1507, “PreNeSt - e:Med”). Author contributions Conceptualization: RGM; Methodology: VB, VK; Data curation: AS; Investigation: RGM; Supervision: RGM; Writing (original draft): RGM, VB, VK; Writing (review and editing): RGM, MDS. Data and materials availability: Raw data analyzed during the current study are not publicly available due to absence of written consent from the participants of the study. Scripts and codes used in the analyses will be deposited in a public database.

---

**Supplementary Figure 1:**
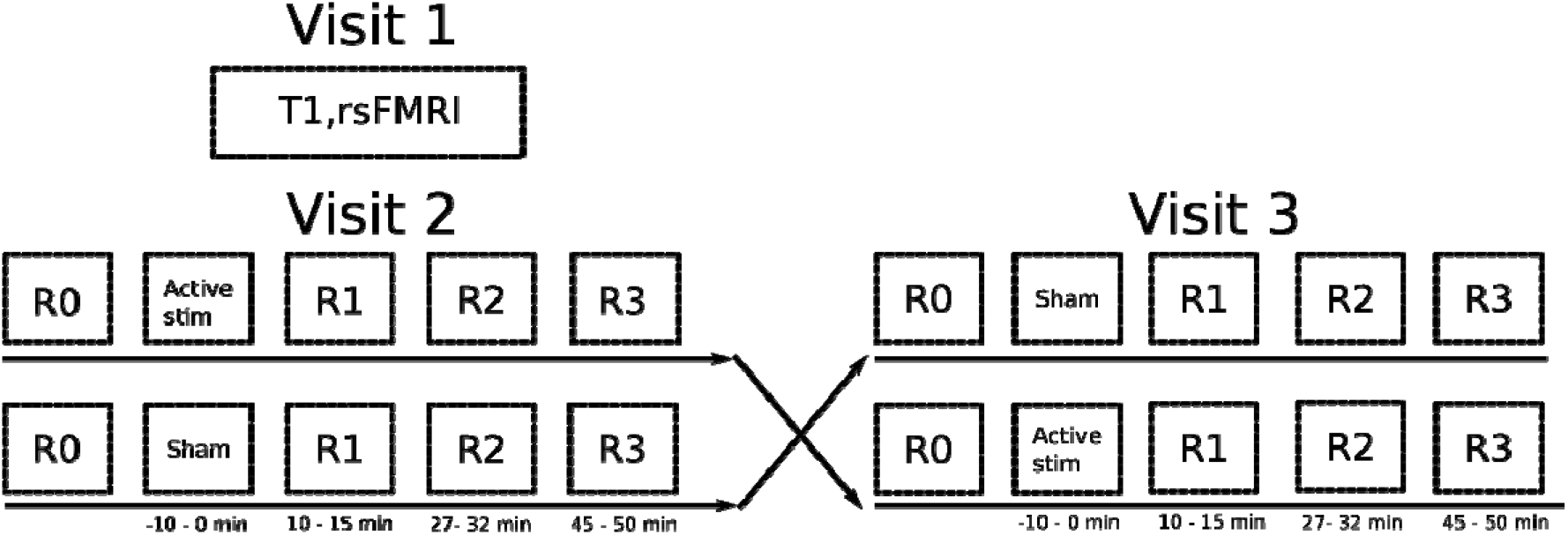
Study design – We acquired T1 and rsfMRI images at visit 1 that were used for personalized target selection. The found target was then located in T1 image for stimulation one week after (Visit 2) and two weeks after (Visit 3) via online neuronavigation. Subjects were assigned to an arm of the study receiving both real and sham in a counterbalanced crossover manner. At the beginning of the sessions on Visit 2 and Visit 3, we obtained a baseline rsfMRI scan (R0). After, the 10 Hz rTMS was delivered at the selected target. Three following scans (R1, R2 R3) were obtained after the stimulation.

**Supplementary Figure 2:**
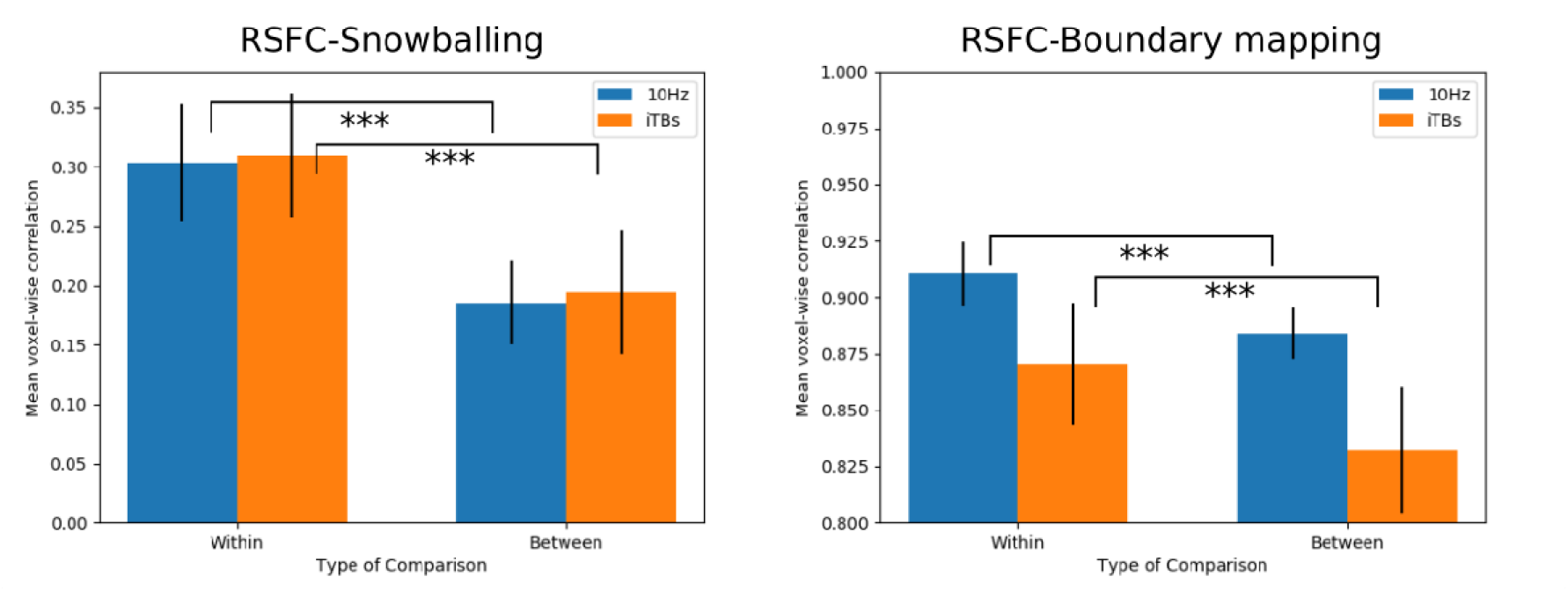
Validation of 3D maps – Spatial correlation within and between node density maps (left) and boundaries (right) in healthy control subjects. Two datasets of baseline rsfMRI separated by about 1 week from independent cohorts of healthy controls (“10Hz” with 23 subjects and “iTBS” with 26 subjects in blue and orange, respectively). For both cohorts, within subject correlation was significantly higher (***p<0.001) than between subjects correlation in both nodes and boundaries. Bars represent standard deviation.

**Supplementary Figure 3:**
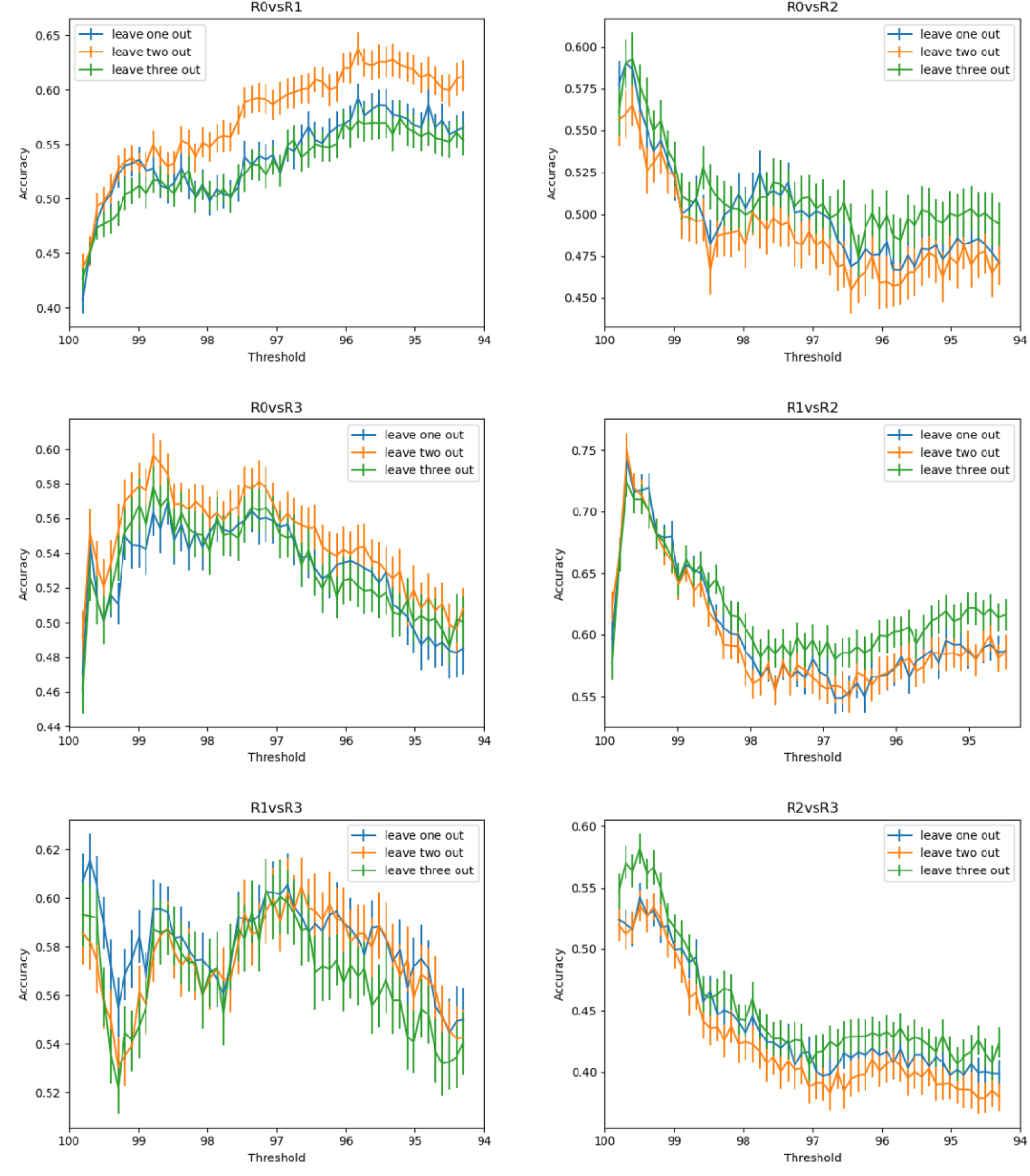
Pairwise classification accuracies of nodes’ density maps using three different cross-validation (CV) strategies

**Supplementary Figure 4:**
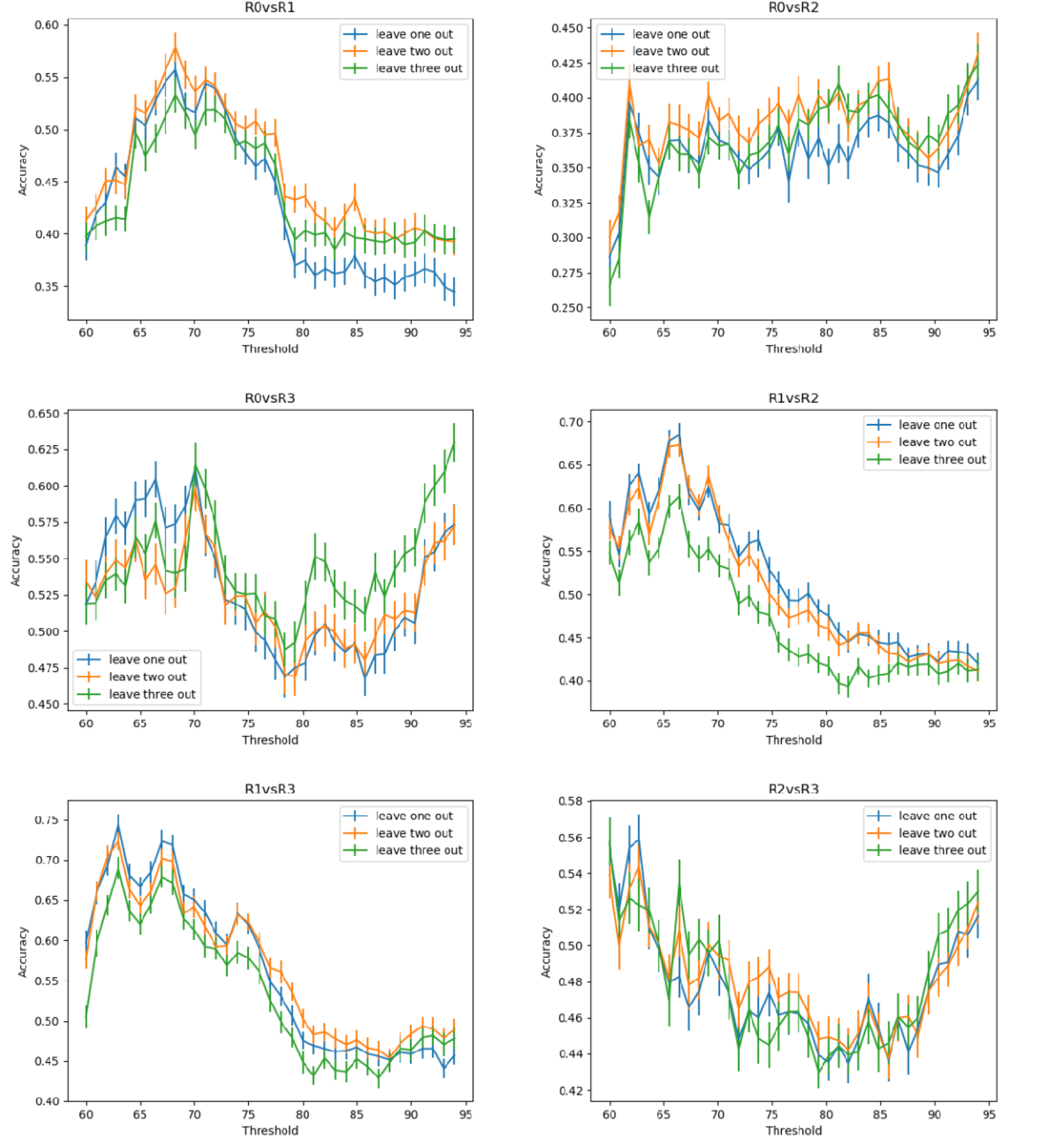
Pairwise classification accuracies of boundary maps using three different cross-validation (CV) strategies

**Supplementary Figure 5:**
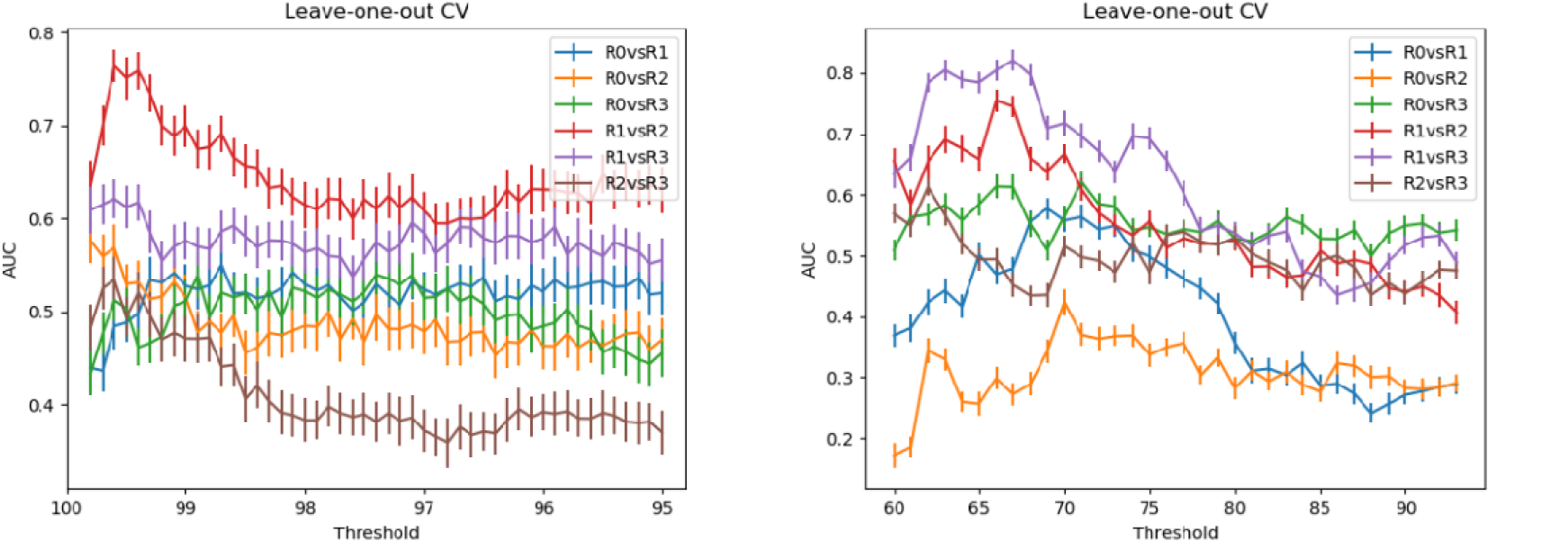
Pairwise classification displayed as area under the curve (AUC) for node (left) and boundary mapping (right)

**Supplementary Table 1:** Highest classification accuracies and area under the curve (AUC) for leave-one-out cross-validation (CV) of nodes and boundaries

|  | Top ranking | Number of voxels | Accuracy (95% CI) [%] | AUC (95% CI) [%] |
| --- | --- | --- | --- | --- |
| **Snowballing peak density** | R1 vs. R2 | 764 | 74.2 (73.0 – 75.4) | 76.4 (74.6 – 78.1) |
| **Boundary mapping** | R1 vs. R2 | 346 | 68.5 (67.2 – 69.8) | 74.5 (72.7 – 86.3) |
|  | R1 vs. R3 | 174 | 74.5 (73.4 – 75.6) | 80.5 (78.7 – 82.2) |

**Supplementary Table 2:** Classification accuracies (%) on head mean frame-to-frame displacement of real and sham rTMS

| Condition | Real<br>Mean (SD) | Sham<br>Mean (SD) | Real - Sham<br>Mean (SD) |
| --- | --- | --- | --- |
| **R0 vs R1** | 0.50 (0.15) | 0.50 (0.29) | 0.57 (0.27) |
| **R0 vs R2** | 0.52 (0.10) | 0.54 (0.20) | 0.52 (0.31) |
| **R0 vs R3** | 0.52 (0.10) | 0.50 (0.00) | 0.35 (0.23) |
| **R1 vs R2** | 0.54 (0.14) | 0.54 (0.20) | 0.46 (0.20) |
| **R1 vs R3** | 0.57 (0.17) | 0.54 (0.14) | 0.52 (0.23) |
| **R2 vs R3** | 0.50 (0.00) | 0.50 (0.00) | 0.50 (0.00) |

